# Functional synchronization between hippocampal sEEG, parietal ECoG and scalp EEG during a verbal working memory task

**DOI:** 10.1101/2020.06.05.136515

**Authors:** Vasileios Dimakopoulos, Ece Boran, Peter Hilfiker, Lennart Stieglitz, Thomas Grunwald, Pierre Mégevand, Johannes Sarnthein

## Abstract

**Background:** The maintenance of items in working memory (WM) relies on a widespread network of brain areas where synchronization between electrophysiological recordings may reflect functional coupling. While the coupling from hippocampus to scalp EEG is well established, we provide here direct cortical recordings for a fine-grained analysis.

**Methods:** A patient performed a WM task where a string of letters was presented all at once, thus separating the encoding period from the maintenance period. We recorded sEEG from the hippocampus, temporo-parietal ECoG from a 64-contact grid electrode, and scalp EEG.

**Results:** Power spectral density (PSD) showed a clear task dependence: PSD in the posterior parietal lobe (10 Hz) and in the hippocampus (20 Hz) peaked towards the end of the maintenance period.

Inter-area synchronization was characterized by the phase locking value (PLV). WM maintenance enhanced PLV between hippocampal sEEG and scalp EEG specifically in the theta range [6 7] Hz.

PLV from hippocampus to parietal cortex increased during maintenance in the [9 10] Hz alpha and the 20 Hz range.

When analyzing the information flow to and from auditory cortex by Granger causality, the flow was from auditory cortex to hippocampus with a peak in the [8 18] Hz range while letters were presented, and this flow was subsequently reversed during maintenance, while letters were maintained in memory.

**Conclusions:** The increased functional interaction between hippocampus and cortex through synchronized oscillatory activity and the directed information flow provide physiological basis for reverberation of memory items during maintenance. This points to a network for working memory that is bound by coherent oscillations involving cortical areas and hippocampus.

**SIGNIFICANCE STATEMENT:** Hippocampal activity is known for its role in cognitive tasks involving episodic memory or spatial navigation, but its role in working memory and its sensitivity to workload is still under debate. Here, we investigated hippocampal and cortical activity while a subject maintained sets of letters in verbal working memory for a few seconds to guide action.

After confirming the coupling between hippocampal oscillations and oscillations on the scalp, we found during maintenance that hippocampal oscillations increased coupling differentially to several areas of cortex by recording directly from the cortex.. During encoding of the letters, information flow was from auditory cortex to hippocampus and subsequently reversed during maintenance, thus providing a physiological basis for memory encoding and maintenance.

This demonstrates a network for working memory that is bound by coherent oscillations that underlie the functional connectivity between cortical areas and hippocampus.

## INTRODUCTION

Working memory (WM) describes our capacity to represent sensory input for prospective use (Baddeley, 2003; Christophel et al., 2017). Maintaining content in WM requires collaboration within a widespread network of brain regions. The anatomical basis of WM was shown noninvasively with EEG / MEG (Sarnthein et al., 1998; Tuladhar et al., 2007; Michels et al., 2008; Näpflin et al., 2008; Polania et al., 2012) and functional magnetic resonance imaging (fMRI) (Wager and Smith, 2003) and invasively with intracranial EEG (Raghavachari et al., 2001; Rizzuto et al., 2003; van Vugt et al., 2010; Maris et al., 2011; Leszczyński et al., 2015; Cogan et al., 2017; Schwiedrzik et al., 2018) and single unit recordings (Kamiński et al., 2017; Kornblith et al., 2017; Boran et al., 2019).

In cortical brain regions, WM maintenance correlates with sustained neuronal oscillations, most frequently reported in the theta-alpha range (3-12 Hz) (Raghavachari et al., 2001; Tuladhar et al., 2007; Michels et al., 2008; Näpflin et al., 2008; Maris et al., 2011; Polania et al., 2012; Hsieh and Ranganath, 2014; Cogan et al., 2017). Also in the hippocampus, WM maintenance was associated with sustained theta-alpha oscillations (van Vugt et al., 2010; Boran et al., 2019). As a hallmark for WM maintenance, persistent neuronal firing was reported during the absence of sensory input, indicating the involvement of the medial temporal lobe in WM (Kaminski et al., 2017; Kornblith et al., 2017; Boran et al., 2019).

At the network level, synchronized oscillations have been proposed as a mechanism for functional interactions between brain regions (Fries, 2015; Pesaran et al., 2018). It is thought that these oscillations show temporal coupling of the low-frequency phase for long-range communication between cortical areas (Sarnthein et al., 1998; Maris et al., 2011; Polania et al., 2012; Solomon et al., 2017; Boran et al., 2019). This synchronization suggests an active maintenance process through reverberating signals between brain regions.

We here extend previous studies with the same task (Michels et al., 2008; Boran et al., 2019), which had shown strong scalp EEG alpha and synchronization between stereotactic EEG (sEEG) in the hippocampus and scalp EEG in 9 subjects. In addition to scalp EEG and hippocampal sEEG, the subject of this study had cortical recordings (ECoG) from an electrode grid spanning the left hemisphere from motor cortex to visual cortex. Given that this implantation scheme is rare for medical reasons, these recordings represent a rare chance. While corroborating the earlier results, the cortical grid allowed for fine-grained analysis of the hippocampal-cortical interaction.

## RESULTS

### Task and behavior

The subject performed a modified Sternberg WM task (2 sessions of 50 trials each). In the task, items were presented all at once rather than sequentially, thus separating the encoding period from the maintenance period. In each trial, the subject was instructed to memorize a set of 4, 6 or 8 letters presented for 2 s (encoding). The number of letters was thus specific for the memory load. The subject read the letters and heard them spoken at the same time. After a delay (maintenance) period of 3 s, a probe letter prompted the subject to retrieve their memory (retrieval) and to indicate by button press (“IN” or “OUT”) whether or not the probe letter was a member of the letter set held in memory (**Fig. 1a**). Throughout this paper, the term “workload” refers to the number of letters held in memory. During the maintenance, the verbal representation of the letter strings is thought to activate verbal WM through the phonological loop with the aim to produce an appropriate behavioral response (Baddeley, 2003).

The average correct response rate was 85% (86% for IN and 73% for OUT trials). The rate of correct responses decreased with set size from a set size of 4 (97% correct responses), to set sizes of 6 (81%) and 8 (72%). The memory capacity was 5.7 (Cowan’s K, (correct IN rate + correct OUT rate −1)*set size), which indicates that the subject was able to maintain at least 4 letters in memory. The mean response time (RT) for correct trials (75 out of 100) was 1.49 ± 0.80 seconds and increased with workload from set size 4 (1.24 ± 0.66 s) to 6 (1.44 ± 0.64 s) and 8 (1.54 ± 0.79 s), 58 ms/item. Correct IN/OUT decisions were made more rapidly than incorrect decisions (1.49 ± 0.80 versus 1.8 ± 0.64 seconds). In summary, these data show that the subject was able to perform the task and that the difficulty of the task increased with set size.

**Figure 1.**
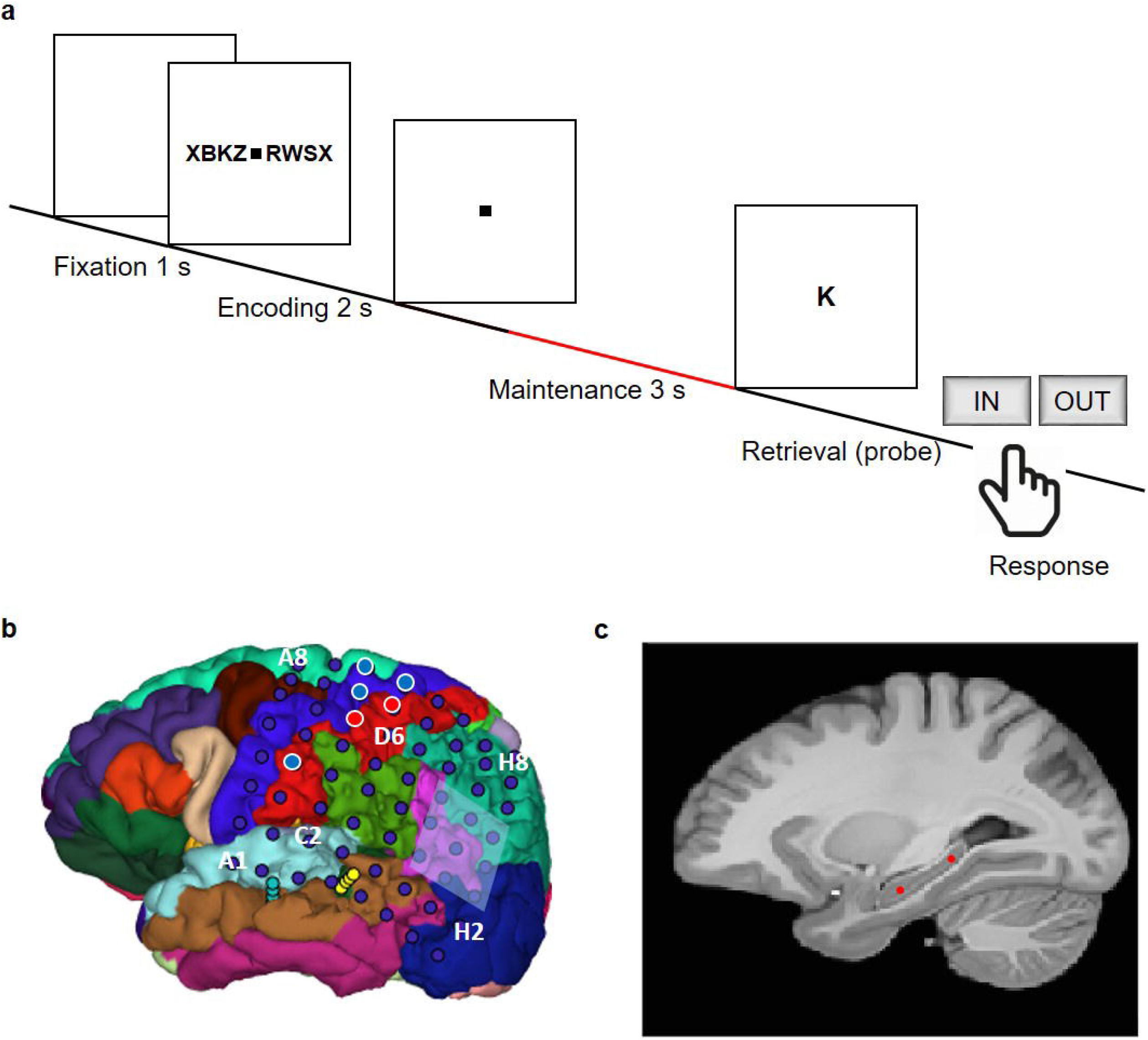
Task and recording sites. (a) In the task, sets of consonants were presented and had to be memorized. The set size (4, 6 or 8 letters) determined WM workload. In each trial, presentation of a letter string (encoding period, 2 s) was followed by a delay (maintenance period, 3 s). After the delay, a probe letter was shown, and subjects indicated whether the probe was presented during the encoding period. (b) Location of the ECoG grid contacts. (c) The tip locations of the hippocampal sEEG electrodes are projected on the parasagittal plane x = −25.2 mm (red markers).

### Spectral power density in cortical and hippocampal recordings

Local synchronization of neuronal activity as characterized by the power spectral density (PSD) of the recordings showed a clear task dependence. In the ECoG of cortical grid electrode (**Fig. 1b**), at contact H3 on the posterior parietal lobe bordering the occipital lobe, the visual presentation of the stimuli during encoding was associated with strong gamma activity (> 40 Hz, **Fig. 2 ab**). During maintenance, the cortical PSD peaked at 12 Hz (**Fig. 2a**) and the relative PSD (**Fig. 2b**) had its maximum towards the end of the maintenance period (**Fig. 2b**). In the sEEG of hippocampus (**Fig. 1c**), the PSD showed a salient peak at 24 Hz during maintenance and a soft peak at 12 Hz during the whole trial (**Fig. 2c**). The relative PSD peaked towards the end of the maintenance period (**Fig. 2d**).

**Figure 2.**
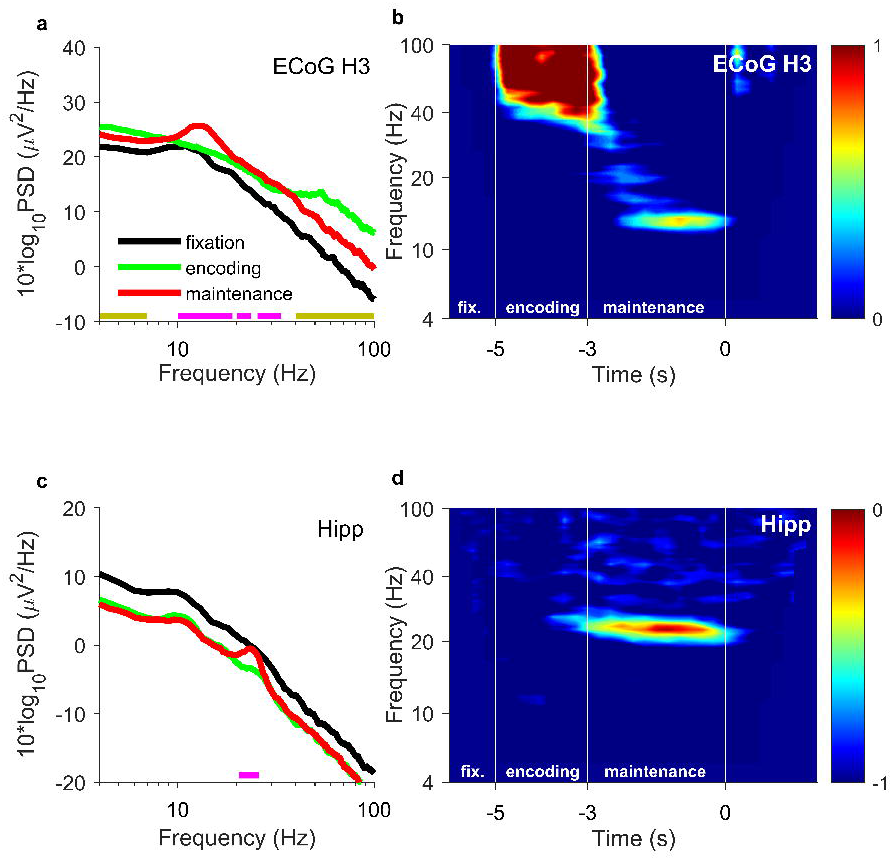
Power Spectral Density (PSD) of hippocampal sEEG and cortical ECoG. a) PSD of ECoG channel H3 over posterior parietal / occipital cortex. During encoding (green), relative PSD is highest in the gamma band (>40 Hz), which reflects visual perception, (golden bar). During maintenance (red), the PSD is highest in the [12 18] Hz band (magenta bar). b) Relative PSD of ECoG contact H3 with respect to the PSD during fixation. The gamma activity (>40 Hz) spans the whole of the encoding period [−5 −3] s and reflects visual perception. The [12 18] Hz activity is strongest towards the end of the maintenance period [−3 0] s towards the presentation of the probe letter at 0 s. c) For hippocampal sEEG, PSD during fixation (black, peak at 10 Hz) is higher than during encoding (green) and during maintenance (red, peaks at 12 Hz and at 24 Hz). d) Relative PSD of the hippocampal sEEG. The [20 26] Hz activity is strongest towards the end of the maintenance period. Bars: significant difference between encoding or maintenance and fixation, P < 0.05, cluster-based nonparametric permutation test. Color scale: relative PSD.

### Cortico-hippocampal functional coupling

To investigate how hippocampal and cortical activities are related, we examined the interregional synchronization with the phase locking value (PLV) as a measure of functional connectivity between different recording sites. In comparisons, we focus on the task conditions fixation and maintenance because the task stimulus was only the fixation square in both conditions (**Fig. 1a**). Between hippocampal sEEG and scalp EEG, the WM maintenance enhanced the PLV specifically in the theta range ([6 7] Hz, to left parietal electrode P3, **Fig. 3a**) and towards the end of the maintenance period (**Fig. 3b**). Among all scalp EEG channels, the PLV was highest to bilateral parietal electrode sites (**Fig. 3c**).

We next investigated the synchronization from scalp electrode P3 to the ECoG grid electrode contacts and found enhanced PLV during maintenance for parietal contact D6 again in the theta range ([6 9] Hz, **Fig. 3d**). The PLV spectra in **Fig. 3ad** showed a distinct theta peak during maintenance. When looking at the theta PLV from P3 to ECoG ([6 9] Hz, **Fig. 3e)**, we find the highest PLV to frontal contact A8. While the PLV to A8 remained constant during the different periods of the task (data not shown) and thereby seems unrelated to the task, it shows that theta PLV between an electrode pair does not decrease as a simple function of distance. To the contrary, the region of high theta PLV from D6 to scalp EEG was widely distributed over ipsilateral and contralateral electrodes (**Fig. 3f**), pointing to a widely distributed theta network.

**Figure 3.**
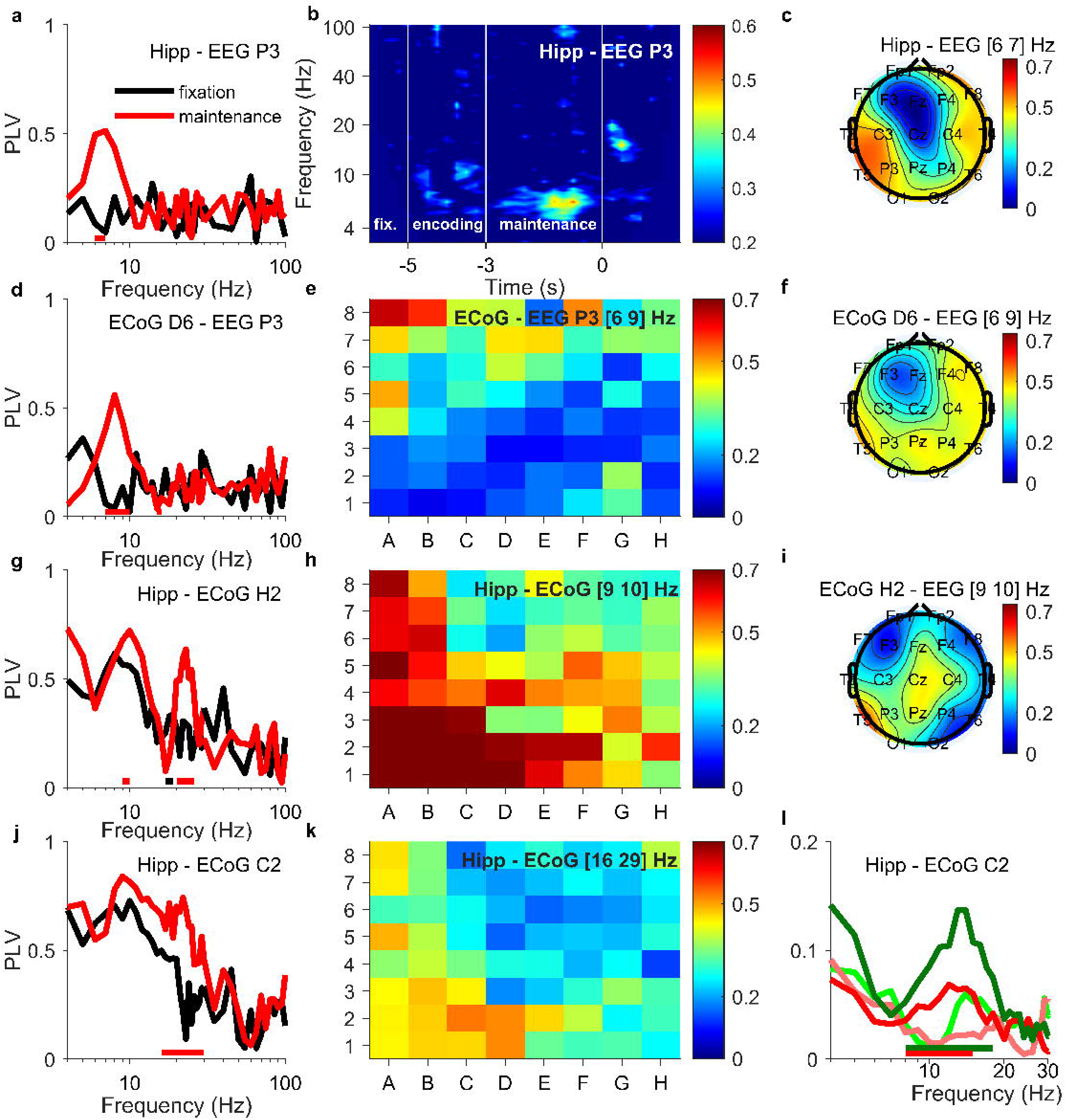
Synchronization between hippocampal sEEG, ECoG and scalp EEG. a) The PLV between hippocampal sEEG and scalp EEG electrode P3 is higher during maintenance (red) than during fixation (black) in the theta band [6 7] Hz. b) In the PLV between hippocampal sEEG and scalp EEG electrode P3 the [6 7] Hz activity is strongest towards the end of the maintenance period [−3 0] s towards the presentation of the probe letter at 0 s. c) During maintenance in the [6 7] Hz theta band, the PLV is high from hippocampal sEEG to electrodes over both left and right parietal cortex. d) Between scalp EEG electrode P3 and parietal ECoG electrode contact D6, PLV during maintenance (red) in the band [6-9] Hz is higher than during fixation (black). e) The PLV in the [6-9] Hz band from P3 is high for ECoG contacts over parietal and motor cortex. f) The PLV between ECoG contact D6 and scalp EEG in the [6-9] Hz band is high to both left and right parietal scalp electrodes. g) Between hippocampal sEEG and ECoG electrode contact H2, PLV during maintenance (red) is higher than during fixation (black) in the bands [9-10] Hz and [20-26] Hz h) The PLV in the [9-10] Hz band from hippocampal sEEG is highest to ECoG contacts over temporal cortex and also high to ECoG contact H2 over posterior parietal / occipital cortex. i) The PLV between ECoG H2 and scalp EEG in the [9 10] Hz band is highest to central scalp electrodes. j) Between hippocampal sEEG and ECoG C2, PLV is high in a wide frequency band. In the band [16-29] Hz, PLV during maintenance (red) is higher than during fixation (black). k) The PLV in the [16-29] Hz band from hippocampal sEEG is highest to ECoG contacts over temporal cortex with a maximum at contact C2 over auditory cortex. l) Spectral Granger causality. During encoding, the signal from ECoG channel C2 over auditory cortex predicted hippocampal sEEG with a peak at 12 Hz ([8 18] Hz, dark green). The dark green curve is higher than the light green curve (information flow from hippocampus to cortex). During maintenance, information flow is reversed: While the cortex-hippocampus information flow is small (light red), hippocampal sEEG predicts ECoG channel C2 ([8 15] Hz, red). Bars: significant difference between maintenance and fixation, P < 0.05, cluster-based nonparametric permutation test. Color scale: PLV

To characterize intracranial functional connectivity at high resolution, we finally turn to the synchronization from the hippocampus to the ECoG grid electrode contacts. PLV to H2 over parietal/occipital cortex increased during maintenance with respect to fixation in the [9 10] Hz alpha and the 20 Hz range (**Fig. 3g**). However, the strongest alpha PLV, which appeared to temporal and frontal contacts (**Fig. 3h**), was not significantly modulated by the task. Finally, PLV to C2 over auditory cortex was enhanced during maintenance (**Fig. 3j**); a robust phenomenon appearing also surrounding contacts (**Fig. 3k**). When analyzing the information flow to and from C2 by Granger causality (**Fig. 3l**), the flow was from auditory cortex to hippocampus in the [8 18] Hz range while letters were presented; this flow was subsequently reversed during maintenance in the [8 15] Hz range, while letters were maintained in memory.

## DISCUSSION

### Windows of activity during the task – “gating”

The subject presented with a particularly clear association of working memory maintenance and enhanced oscillatory brain activity. The effect was found both in hippocampus and cortex (**Fig. 2**), and in the functional coupling between the two (**Fig. 3**). The selective activity increase during maintenance has been termed “gating” (Raghavachari et al., 2001) and varies in strength across subjects for the same task (Michels et al., 2008; Näpflin et al., 2008; Boran et al., 2019; Boran et al., 2020). The separation of WM trial periods was expected, as single-neuron data also showed period-dependent differences in WM activity (Kamiński et al., 2017; Kornblith et al., 2017; Boran et al., 2019) during object maintenance.

The evolution over time shows the activity to increase during the last 2 s of the maintenance period and to rapidly diminish following the probe letter (**Fig. 2bd, 3b**). For these sites, neural communication between brain regions was channeled through coupling within a limited time window between the onset and offset of the maintenance phase. This fine-grained time course suggests that the WM network function goes beyond memorization toward a task-related preparation in expectance of the probe.

### Maintenance-induced connectivity to parietal EEG

We first discuss how the intracranial recording sites sEEG and ECoG communicate with scalp EEG within the WM network. The data from this subject corroborates a result from an earlier study (Boran et al., 2019) that maintenance consistently elicited enhanced theta PLV (**Fig. 3ab**) to parietal scalp EEG (**Fig. 3c**). Similarly, several ECoG sites showed task-related PLV synchronization to parietal scalp EEG (**Fig. 3def**). This finding is consistent with parietal scalp EEG being a common locus for alpha waves and findings in several healthy subjects performing the same (Michels et al., 2008; Näpflin et al., 2008) or similar tasks (Tuladhar et al., 2007). In agreement with the stimulus material of the task, letter stimuli elicit persistent activity at parietal cortical sites (Christophel et al., 2017).

### Maintenance-induced connectivity between hippocampal sEEG and ECoG

The wide distribution of recording sites on the ECoG grid enables a fine-grained spatial sampling of cortical activity. To rule out non-functional high PLV due to volume conduction, we view as task-related only those findings that show a significant difference between the task conditions fixation and maintenance because in both conditions only the fixation square was present as a stimulus.

A large number of ECoG contacts showed enhanced synchronization to hippocampus during maintenance (**Fig. 3**), among them contacts over auditory cortex (**Fig. 3k**) and contacts over visual cortex (**Fig. 3h**). The latter site coincides with the cortical generator of scalp EEG that was found in the cuneus for the same task (Michels et al., 2008). In some hippocampal-cortical contact pairs, PLV reached significance both around 10 Hz and around 20 Hz, e.g. H2 (**Fig. 3g**); reminiscent of the 12 Hz and 24 Hz peak in the hippocampal PSD during maintenance (**Fig. 2c**). We view the two frequencies as harmonics of the same physiological generator that is active during working memory maintenance. The long-range functional synchronization between hippocampal and cortical sites would then resemble cortical findings during WM tasks (Sarnthein et al., 1998; Polania et al., 2012) and other tasks (Solomon et al., 2017) that activate oscillations in long-range recurrent connections (Fries, 2015; Pesaran et al., 2018).

### Connectivity during encoding and maintenance

The connectivity from hippocampus to ECoG over auditory cortex stood out. Already during encoding there was high PLV and the analysis of Granger causality indicated information flow from auditory cortex to hippocampus while the subject heard the letters (**Fig. 3l**). This would match the encoding of auditory information in a brain network that involves neuronal firing in the hippocampus (Boran et al., 2019). During maintenance, this information flow was reversed, i.e. from hippocampus to cortex. This points to the involvement of hippocampus in the maintenance of items in working memory (Jeneson and Squire, 2012). The involvement of auditory cortex is plausible because the verbal representation of the letter strings is thought to activate verbal WM through the phonological loop (Baddeley, 2003; Christophel et al., 2017).

## CONCLUSIONS

While the subject performed the task, we observed increased functional interaction between hippocampus and cortex through synchronized oscillatory activity. This corroborates earlier results with the same task in nine patients. In addition, the cortical grid electrode allowed a more fine-grained analysis. During encoding of the letters, information flow was from auditory cortex to hippocampus and subsequently reversed during maintenance, thus providing a physiological basis for reverberation of memory items. This points to a network for working memory that is bound by coherent oscillations involving cortical areas and hippocampus.

## METHODS

### Task

We used a modified Sternberg task in which the encoding of memory items, maintenance, and retrieval were temporally separated (**Fig. 1a**). Each trial started with a fixation period ([−6, −5] s), followed by the stimulus ([−5, −3] s). The stimulus consisted of a set of eight consonants at the center of the screen. The middle four, six, or eight letters were the memory items, which determined the set size for the trial (4, 6, or 8 respectively). The outer positions were filled with “X,” which was never a memory item. The subject read the letters and heard them spoken at the same time. After the stimulus, the letters disappeared from the screen, and the maintenance interval started ([−3, 0] s). A fixation square was shown throughout fixation, encoding, and maintenance. After maintenance, a probe was presented. The subject responded with a button press to indicate whether the probe was part of the stimulus. The subject was instructed to respond as rapidly as possible without making errors. The subject was right handed and used the right hand for the response. After the response, the probe was turned off, and the subject received acoustic feedback regarding whether the response was correct or incorrect. The subject performed two sessions of 50 trials, which lasted approximately 10 min each. Trials with different set sizes were presented in a random order, with the single exception that a trial with an incorrect response was always followed by a trial with a set size of 4. The task is freely available at www.neurobs.com/ex_files/expt_view?id=266.

### Subject and electrodes

One subject (46y, f) participated in the study. The patient had a medically intractable focal epilepsy in the presence of both a traumatic brain injury in the left parietal lobe acquired during childhood and left-sided hippocampal sclerosis. To investigate a potential surgical treatment of epilepsy, the patient was implanted with intracranial electrodes. The subject provided written informed consent for the study, which was approved by the institutional ethics review board (PB 2016-02055). The subject had corrected-to-normal vision and was right handed as confirmed by neuropsychological testing. The depth electrodes (1.3 mm diameter, 8 contacts of 1.6 mm length, spacing between contact centers 5 mm, ADTech®, Racine, WI, www.adtechmedical.com) were stereotactically implanted into the hippocampus. The ECoG was collected using an 8 x 8 electrode grid (Ad – Tech, Racine, WI) placed directly on the cortex. Platinum electrodes with 4 mm^2^ contact surface and 1 cm inter-electrode distances were used. In addition, scalp EEG electrodes were placed at the sites of the 10-20 system with minor adaptations to avoid surgical scalp lesions.

### Electrode localization

To localize the ECoG grid, we used the subject’s postoperative MR, aligned to CT and produced a 3D reconstruction of the subject’s pial brain surface. Grid electrode coordinates were projected on the pial surface as described in (Groppe et al., 2017) (**Fig. 1b**). The stereotactic sEEG electrodes were localized using post-implantation computed tomography (CT) and post-implantation structural T1-weighted MRI scans. The CT scan was registered to the post-implantation scan as implemented in FieldTrip (Stolk et al., 2017). A fused image of CT and MRI scans was produced and the electrode contacts were marked visually. The contact positions were projected on the patient’s MRI (**Fig. 1c**).

### Recording setup and re-referencing

All recordings were performed with Neuralynx ATLAS, sampling rate 4000 Hz, 0.5-1000 Hz passband (Neuralynx, Bozeman MT, USA, www.neuralynx.com). ECoG and sEEG were recorded against a common intracranial reference and then re-referenced against a sEEG contact in the white matter. As hippocampal signal (Hipp), we selected the contact 2 of the anterior hippocampal sEEG electrode, which was localized in the hippocampal head. The scalp EEG was recorded against an electrode near the vertex and was then re-referenced to the averaged mastoid channels.

### Data preprocessing

All signals were downsampled to 500 Hz. Trials with large unitary artefacts in the scalp EEG were rejected. The scalp EEG was then decomposed using Independent Component Analysis (ICA) using FieldTrip (Oostenveld et al., 2011). Components with eye movement artefacts were rejected and the EEG reconstructed. We selected only trials with good data quality, correct response and high workload for further analysis (set sizes 6 and 8, 43/100 trials).

### Power spectral density (PSD)

To calculate the power spectral density (PSD) (**Fig. 2**), we transformed the data to the frequency domain using the multitaper method based on discrete prolate spheroidal sequences (2 tapers, frequency range 4 to 100 Hz with frequency resolution of 0.5 Hz, ± 2 Hz smoothing) as implemented in FieldTrip to reduce spectral leakage and control the frequency smoothing. The multitaper method modifies K periodograms obtained from different spheroidal sequences as the window.

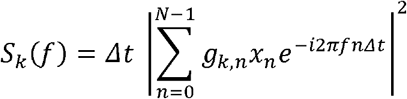

where *S_k_*(*f*) denotes the modified periodogram using the k-th spheroidal sequence *g_k,n_*, of length n, *x_n_* the data at the window n with sampling interval *Δt* Afterwards the multitaper method averages the K modified periodograms to calculate the PSD.

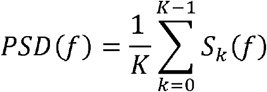

### Phase-locking value (PLV)

To evaluate the functional connectivity among hippocampus, cortex and other underlying structures we calculated the phase-locking value (PLV) between scalp EEG and ECoG grid, scalp-EEG and sEEG channels on hippocampus but also between sEEG channels and ECoG grid (multitaper frequency transformation with 2 tapers based on Fourier transform, frequency range 4-100 Hz with frequency resolution of 1 Hz).

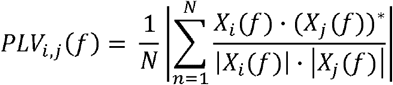

where PLV_i,j_ is the PLV between channels i,j, N is the number of trials, X(f) is the Fourier transform of x(t), and (·)* represents the complex conjugate.

Using the spectra of the two-second epochs, phase differences were calculated for each electrode pair (i,j) to quantify the inter-electrode low-frequency phase coupling. The phase difference between the two signals indexes the coherence between each electrode pair and is expressed as the PLV. The PLV ranges between 0 and 1, with values approaching 1 if the two signals show a constant phase relationship over all trials.

### Time-frequency analysis

To calculate the PSD in the time-frequency domain (**Fig. 2bd**), each trial was transformed to a time-frequency map. We used multitapers with a window width of 10 cycles per frequency point, smoothed with 0.2 × frequency, and used three tapers. We computed power in the frequency range of 4 to 100 Hz with a time resolution of 0.1 s. The PSD during fixation ([−6.0, −5.0] s) served as a baseline for the baseline correction (PSD(t) − PSD(fixation))/ PSD(fixation) for each time-frequency point. Using the same parameters, we computed the PLV in the time-frequency domain (**Fig 3b**).

### Spectral Granger causality analysis

In order to evaluate the direction of information flow between the hippocampus and the cortex, we calculated spectral non-parametric Granger causality (GC) for pairs of ECoG grid and sEEG hippocampus channels during different task periods (**Fig.3l**). GC examines if the activity on one channel can forecast activity in the target channel. In the spectral domain, GC measures the fraction of the total power that is contributed by the source to the target. We transformed signals to the frequency domain using the multitaper method in the same way as for the spectral power. We used a non-parametric spectral approach to measure the interaction in the channel pairs at a given interval time (Bastos and Schoffelen, 2016). In this approach, the spectral transfer matrix is obtained from the Fourier transform of the data. We used the FieldTrip toolbox to factorize the transfer function H(f) and the noise covariance matrix Σ. The transfer function and the noise covariance matrix was then employed to calculate the total and the intrinsic power, S(f) = H(f)ΣH*(f), through which we calculated the Granger interaction in terms of power fractions contributed from the source to the target.

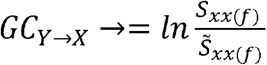

where *S_xx(f)_* is the total power and 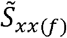 the instantaneous power.

### Statistics

For statistical significance analysis we used cluster-based nonparametric permutation tests. To assess the significance of PLV values between a pair of channels during a specific task period, we compared the true values to a null distribution. We recomputed PLV after shuffling the trial number for a single channel in the pair, while keeping the trial number of the other channel constant. We repeated this n = 200 times to create a null distribution of PLV. The null distribution was exploited to calculate the percentile threshold P = 0.05. We repeated the same procedure for the assessment of the significance of Granger causality spectra.

To assess the significance of the difference of the PLV between different task periods, we compared the difference of the true values to a null distribution of differences. We recomputed PLV after switching task periods randomly across trials, while keeping the trial numbers for both channels constant. Then we computed the difference of PLV for the two conditions compared. We repeated this n = 200 times to create a null distribution of differences. The null distribution was exploited to calculate the percentile threshold P = 0.05. We repeated a similar procedure for assessing the significance of the difference of PSD between different task periods. Only the frequency bins identified above the threshold were marked as statistically significant.

## Acknowledgments

We thank the physicians and the staff at Schweizerische Epilepsie-Klinik for their assistance and the patients for their participation. We acknowledge grants awarded by the Swiss National Science Foundation (SNSF 320030_176222 to J. S.) and SNSF Ambizione fellowship (PZ00P3_167836 to P. M.). The funders had no role in the design or analysis of the study.

## Author contributions

J.S. designed the experiments. P.H. set up the recordings. J.S. performed the experiments. V.D, E.B., and J.S. analyzed the data. T.G. provided patient care. L.S. performed surgery. P.M. visualized the grid electrode position, V.D., E.B, and J.S. wrote the manuscript. All of the authors reviewed the final version of the manuscript.

## Ethical considerations

The subject provided written informed consent for the study, which was approved by the institutional ethics review board (PB 2016-02055).

## Data availability

All data needed to evaluate the conclusions in the paper are present in the paper. Additional data related to this paper may be requested from the authors.

